# A flexible quality metric for electrophysiological recordings across brain regions and species

**DOI:** 10.64898/2026.03.06.710130

**Authors:** Noam Roth, Gaelle Chapuis, Olivier Winter, International Brain Laboratory, Ryan A. Ressmeyer, Luke M. Bun, Ryan A. Canfield, Gregory D. Horwitz, Nicholas A. Steinmetz

## Abstract

The increasing size of electrophysiological datasets has heightened the need for quality metrics that automatically reject neurons whose activity was recorded with low sensitivity or specificity. One key approach estimates artifactual contamination by assuming that each neuron has a refractory period (RP), a brief time interval following each action potential when further activity cannot occur. However, existing methods cannot be applied without prior knowledge of the neurons’ RP durations, limiting their usefulness in datasets that include neurons from brain regions or species in which RP durations have not been systematically characterized. Here, we find that neurons in some brain regions (thalamus) and species (macaque) have shorter RP durations than commonly assumed, and we introduce a new metric, the *Sliding Refractory Period* metric, which is robust to variation in a neuron’s RP duration without tuning. We validate the method using simulations, demonstrating that it improves acceptance of uncontaminated spike trains with short or long RP durations while still rejecting contaminated ones. Moreover, by incorporating Poisson statistics into the calculation, the method also improves on prior work by allowing the user to approximately control the false acceptance rate. Our new metric improves quantification of contamination in electrophysiological recordings and enables application of a single tuning-free quality metric to data recorded from diverse brain regions and species.

## Introduction

Technologies that measure activity of individual neurons by the hundreds or thousands have ushered the scientific study of the brain into a new era (Steinmetz et al., 2018; International Brain Lab et al., 2025). A critical challenge for the reproducible analysis of such large-scale physiological datasets is the development of quality metrics that can be applied automatically to flag or discard corrupted recordings of neurons. Ideal metrics should be applicable to a wide variety of datasets without requiring detailed manual tuning in order to maximize reproducibility and interpretability.

For electrophysiological measurements, data corruption can arise due to the presence of noise, the inability to distinguish activity from neighboring neurons, or the physical movement of neurons relative to the measurement device (*drift*), among other factors. In the *spike sorting* process, algorithms attempt to automatically detect action potential events, or *spikes*, and cluster them into *units*, which are collections of spikes with similar waveforms (Quiroga, 2012; Buccino et al., 2022). Ideally, each unit includes all and only spikes from a single neuron. However, errors can occur. On the one hand, spikes of the neuron may be undetected or assigned to other units (*missing spikes*). On the other hand, spikes due to noise or produced by other neurons (*contaminating spikes*) may be incorrectly assigned to the unit corresponding to the neuron under investigation (the *base neuron*). Missing spike rates can be estimated by examining the amplitudes or shapes of assigned spikes for evidence of a detection threshold or classification boundary across which spikes may have been lost (Hill et al., 2011; International Brain Laboratory, 2022; Fabre et al., 2023), or by examining spike rates for anomalous periods of silence (*presence ratio*, (Buccino et al., 2020)). Contamination rates can be estimated by examining the intervals between spikes for instances in which the base neuron’s refractory period has been violated (Hill et al., 2011; Llobet et al., 2022). A final class of quality metric attempts to qualitatively or quantitatively estimate the likelihood of either kind of error by assessing the uniqueness of the waveforms of assigned spikes relative to neighbors (*isolation distance* and related approaches (Schmitzer-Torbert et al., 2005; Chung et al., 2017)).

Here we focus on estimation of contamination by examination of refractory period violations. Prior approaches to this problem start by assuming a certain refractory period (RP) duration, typically 2 or 3 ms. They then quantify the number of spikes observed with shorter inter-spike intervals (ISIs) than this duration, and finally calculate an estimated total contamination rate from the assumptions that these short-ISI spikes reflect contamination and that the contaminating spikes occur at a fixed rate with Poisson occurrence (Hill et al., 2011; Llobet et al., 2022). If the RP duration is mis-specified for a given unit, then this calculation will return biased results: if the base neuron’s true RP duration is shorter than assumed, then real spike pairs will be interpreted as reflecting contamination by the algorithm. It may therefore incorrectly report a higher contamination rate than true one. This type of error is a conservative one, resulting in rejection of uncontaminated units but not resulting in acceptance of contaminated ones. However, if this type of error applies to some neuron populations differently than others, it will introduce a bias in the populations of neurons that are accepted for analysis. And more generally, if the RP duration parameter is badly mis-specified, as when the true RP durations of the neurons in a population are mostly or entirely lower than the assumed duration, then this type of error will result in a decimation of the population of neurons available for analysis. A more subtle type of inaccuracy can occur when the RP duration is mis-specified in the opposite direction: if the true RP duration is longer than assumed, then the calculation will be performed with only a subset of the data that should be available to it. For example, if 2 ms is assumed but the neuron’s true RP duration is 5 ms, only spikes with ISI less than 2 ms will count as contamination when spikes up to ISI of 5 ms ought to have counted. Since only very brief windows are available for estimating these contaminating spikes, the procedure is noisy and generally underpowered, so its performance could be substantially improved by the availability of this extra data.

The validity of the assumed RP duration is therefore critical to obtain accurate results from existing methods. However, RP durations have not been systematically quantified in most brain regions and species (Kara et al., 2000; Bar-Gad et al., 2001), let alone any of the thousands of genetically distinct cell types in the mammalian brain (Yao et al., 2023). Researchers must typically examine their own data manually to estimate the RP durations of neurons they record, a process that is laborious, prone to error, and impractical for datasets with many brain regions or neuron types. Moreover, even if a characteristic RP duration could be established for a specific brain region, this would not account for variability in this parameter from neuron to neuron. As a specific example, parvalbumin-expressing basket cells, a common cortical interneuron, are known to have a fast-spiking phenotype characterized by short RP durations relative to neighboring pyramidal neurons (Hu et al., 2018), so for any cortical recording, a single assumed RP duration will never be equally valid for all types of neurons that might be sampled. In order to maximize effectiveness of this quality metric, it is therefore imperative to have a metric which is robust to differences in RP duration across neurons.

A second major limitation of prior approaches is that they compute only a point estimate of the contamination rate, without accounting for the statistical variability inherent in the observation. Because spikes are generated by stochastic processes, the number of RP violations observed in a finite recording is itself a random variable: a unit with 10% contamination might, by chance, produce few or no violations. This can be especially likely when the recording duration is brief or the firing rate is low, providing few opportunities to observe such contaminating spikes. Prior approaches are therefore significantly biased as a function of firing rate (Vincent and Economo, 2024). In the extreme case, prior methods will estimate 0% contamination in cases with zero observed violations and these units will pass the test, regardless of how statistically underpowered the observation was. This means the false acceptance rate of existing methods is uncontrolled and varies with firing rate and recording duration, growing arbitrarily large when statistical power is low. A more principled approach would compare the observed violation count to the distribution of counts expected under a given contamination level, allowing the user to specify and control the acceptable probability of incorrectly accepting a contaminated unit.

Here we introduce a spike sorting quality metric, the “Sliding RP” metric, which avoids any assumption about RP duration and incorporates Poisson statistics to approximately control false acceptance rates. Motivating this method, we compare estimated RP durations across three brain regions (isocortex, hippocampus, and thalamus in mouse) and across two species (mouse and macaque) revealing substantial differences between regions and species. With simulated data, we validate the performance of the algorithm across a range of realistic parameters. We compare performance to prior state of the art, demonstrating how our approach improves performance on key test cases and enhances interpretability. We provide implementations of the metric in MATLAB and in Python, and within the SpikeInterface package (Buccino et al., 2020).

## Results

### Refractory period (RP) durations vary across brain regions and species

To determine the extent to which RP durations vary across brain regions and species, we analyzed five datasets, and observed a wide range of apparent RP durations (Fig. 1). Three of the datasets are previously-published sets of recordings from mice which each include neurons from visual cortex, hippocampus, and thalamus (Steinmetz et al., 2019; Siegle et al., 2021; International Brain Laboratory et al., 2025). The other two are sets of recordings from macaque lateral geniculate thalamus (LGN; (Ressmeyer et al., 2026)) and unpublished sets of recordings from macaque visual cortex. For the units in each dataset, we estimated the refractory period by fitting a constrained sigmoid function to each unit’s autocorrelogram and extracting the time at which the sigmoid rose above a threshold (Fig 1b; see Methods). We found that refractory periods were significantly shorter in thalamus than in cortex and hippocampus in the mouse (p<0.001, ANOVA, effect of brain region), and significantly shorter in visual cortex of the monkey than the mouse (p<0.001, ANOVA, effect of species; Fig 1c). Importantly, many RP durations in each dataset were shorter than 2 ms, and RP durations varied substantially within each dataset and brain region (Figure 1a,b; S1a). The results of this analysis did not differ significantly when contaminated units were excluded (using the Sliding RP method described below; Figure S1b).

**Figure 1.**
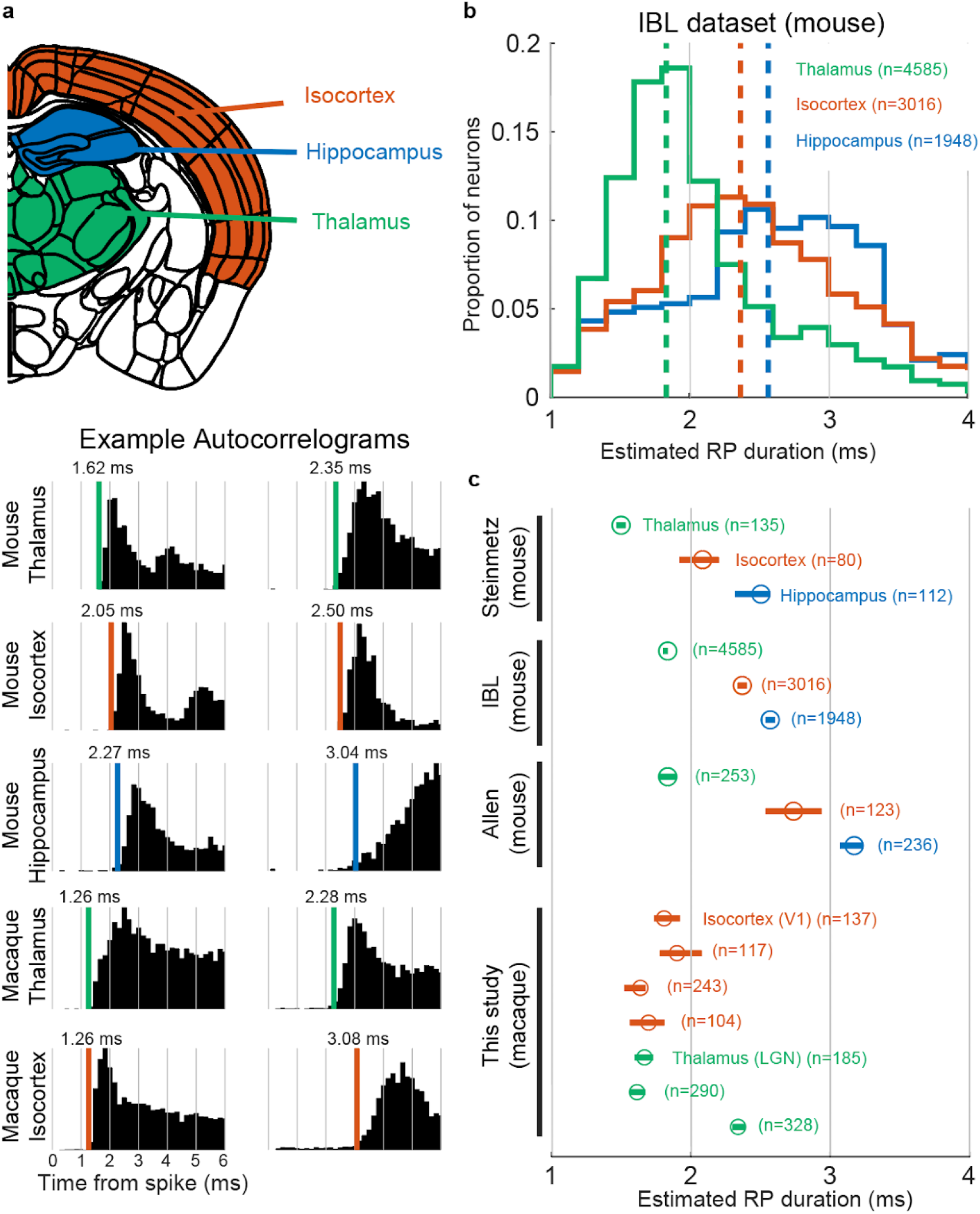
Refractory periods vary across brain regions and species. **a**, Top: coronal slice of the mouse brain atlas (Allen Common Coordinate Framework (Wang et al., 2020)) illustrating the three regions compared: isocortex (orange), hippocampus (blue), and thalamus (green). Bottom: example autocorrelograms from individual units in each region and species, with estimated refractory period duration indicated by the colored vertical line. Two examples are shown per region, illustrating the range of RP durations observed. **b**, Distribution of estimated RP durations for units recorded from thalamus, isocortex, and hippocampus from the IBL dataset (mouse). Dashed vertical lines indicate medians. **c**, Median estimated RP duration by brain region across datasets. Error bars indicate bootstrapped 95% confidence intervals of the median.

### A quality metric suitable for variable RP durations

Given the variability we observed across brain regions and species, we sought to develop a quality metric that could capture the contamination of a unit without prior knowledge of the base neuron’s true RP duration. We first briefly describe the prior state-of-the-art metric, then give an overview of the Sliding RP algorithm (Fig. 2). After introducing the method, we quantify the performance of the method relative to prior state of the art on simulated data (Fig. 3).

**Figure 2.**
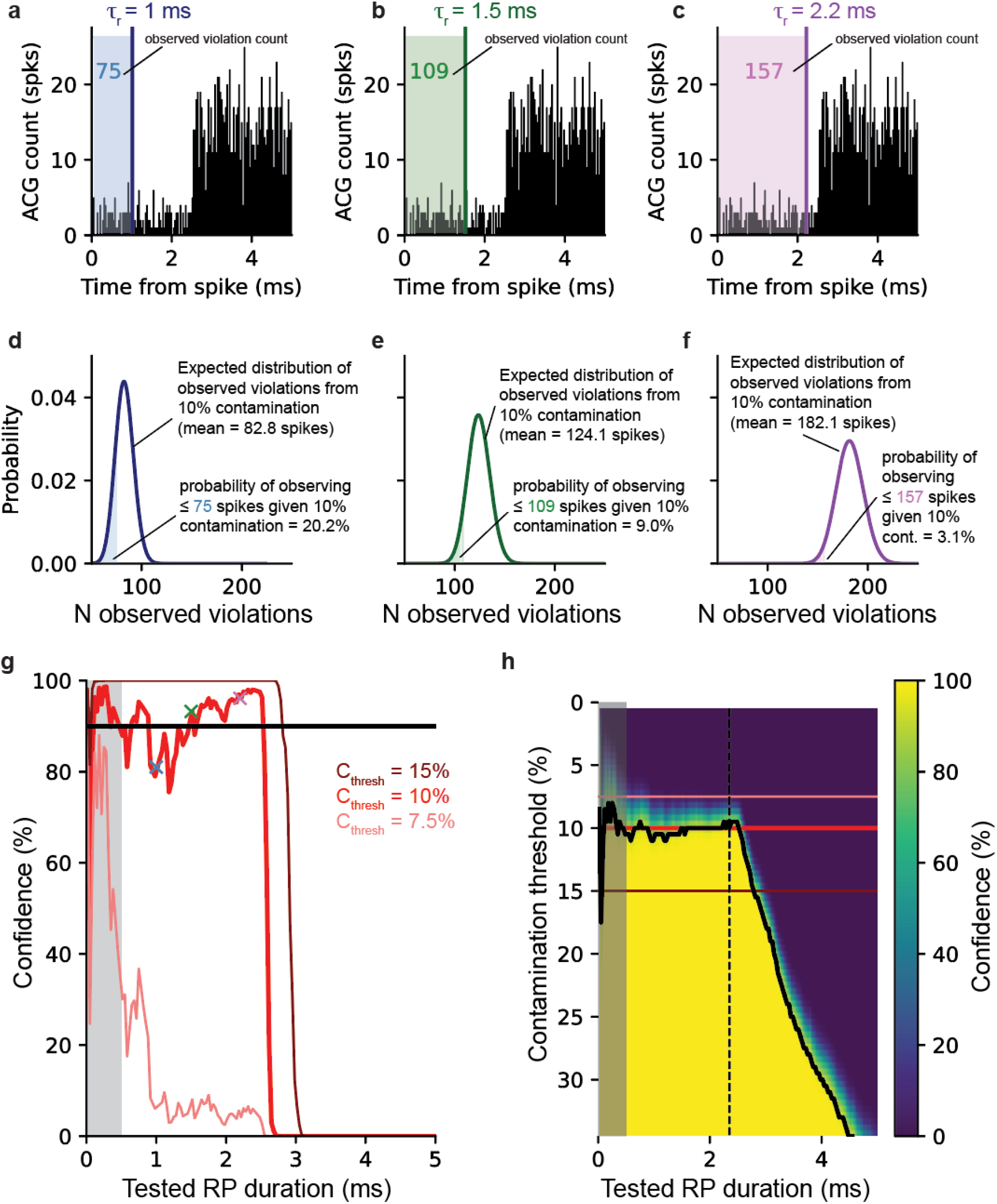
Illustration of the Sliding RP algorithm. **a**, Autocorrelogram of a spike train simulated to be Poisson except for a RP of 2.5 ms duration, with 10% contamination. Here **τ**_r_, the present value of the sliding analysis window, is 1ms, indicated by dark blue vertical line. The count of spikes occurring with shorter intervals than **τ**_r_ is indicated in the shaded region. **b**, Same as in a but for **τ**_r_ = 1.5 ms. **c**, Same as in a but for **τ**_r_ = 2.2 ms. **d**, Confidence computation corresponding to **a**. A Poisson distribution whose mean is the expected number of violations for the example spike train (from a) shown under the assumption that the unit has 10% contamination. The shaded area corresponds to the probability of observing less than or equal to the actual observed number of violation spikes (75, in this case) if the true contamination level were 10%. The *confidence* that contamination is less than or equal to that contamination level is therefore computed as the complement of this probability, indicated by the correspondingly colored x in g. **e**, Confidence computation corresponding to **b**, labelling conventions as in **d. f**, Confidence computation corresponding to c, labelling conventions as in d. **g**, Confidence scores for all possible **τ**_r_, computed as in d, e, and f for a level of 10% contamination (thick red line). Values shown in examples d-f are indicated by the colored Xs. Confidence scores for levels of 7.5% (light red) and 15% (dark red) contamination thresholds (C_thresh_) are also shown. Grey shaded bar indicates time points excluded from the test. 90% confidence level indicated by the horizontal black line. **h**, Full confidence matrix for this spike train, for all tested values of C_thresh_ (y-axis). Confidence is indicated by the color bar. Confidence values shown in g are indicated by the color values along the red horizontal lines corresponding to 7.5%, 10%, and 15% contamination thresholds. Black isocurve indicates 90% confidence. The time point showing minimum contamination with greater than 90% confidence is indicated by the dashed vertical line.

**Figure 3.**
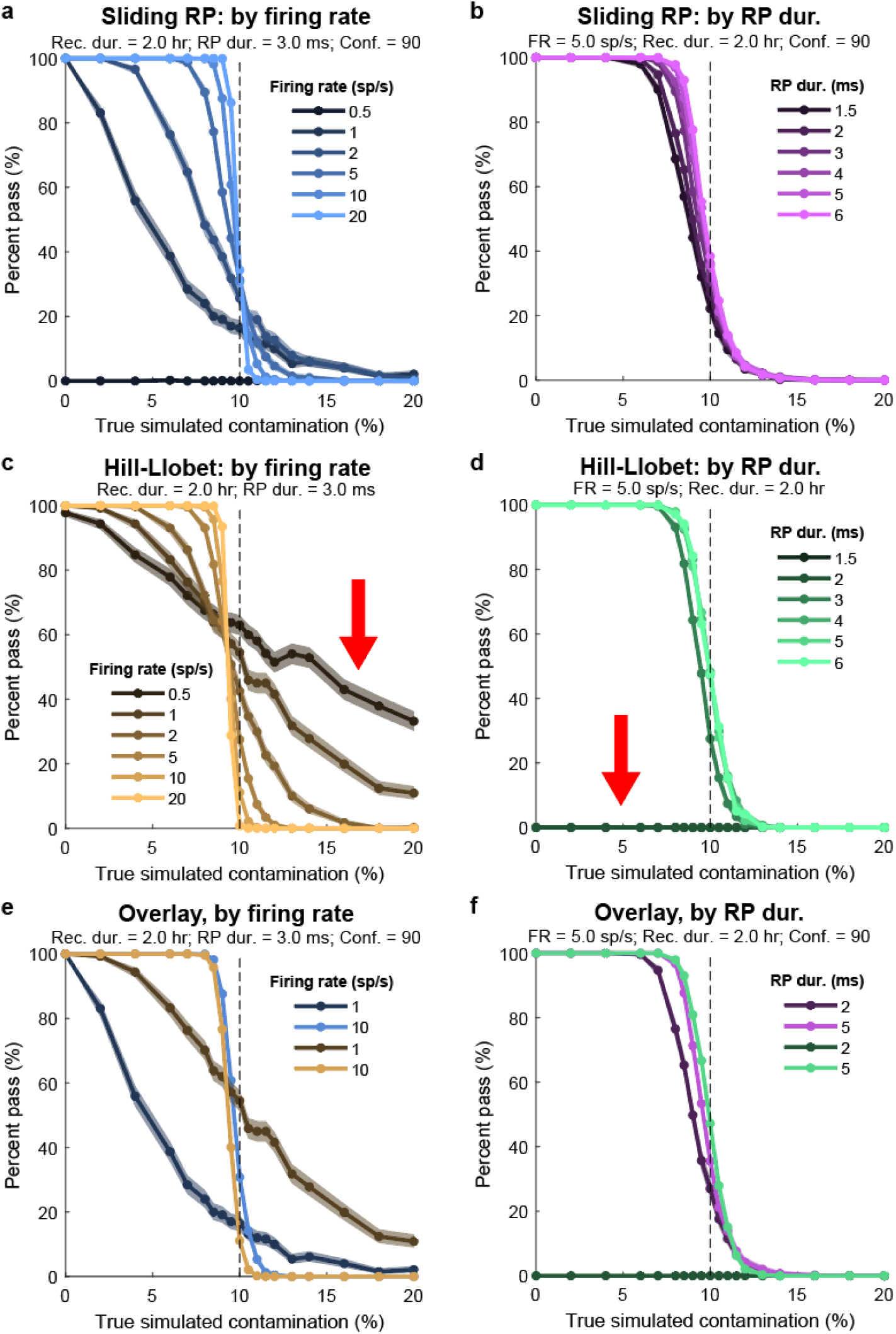
Sliding RP algorithm performance on simulated data, and comparison to prior state of the art. **a**, Percentage of simulated spike trains passing the test as a function of true simulated contamination, for varying firing rates. The acceptable contamination threshold was set at 10% (vertical dashed line). n=1000 simulations per condition; shaded regions reflect 95% confidence intervals (Clopper-Pearson). Other simulation parameters indicated in plot title (“Rec. dur.”, recording duration; “RP dur.”, refractory period duration; “Conf.”, confidence threshold; “FR”, firing rate). **b**, Same as a, varying true RP duration of the simulated spike train. **c-d**, Same as a-b, using the Hill-Llobet method with a fixed RP duration parameter of 3 ms. Red arrow in c points out the high rate of passing contaminated spike trains. Red arrow in d points out the rejection of uncontaminated spike trains when their RP durations were too low. **e**, Overlay of selected traces from panels a and c, with colors as in those panels. **f**, Overlay for selected RP durations.

The aim of the metric is to estimate how much of a spike train consists of contaminating spikes, i.e. those arising from any neurons or noise sources other than the base neuron in question. Any inter-spike intervals (ISIs) that are too brief to be physiologically possible are considered violations of the RP, and the number of these violations can be used to infer the total proportion of the spike train that consists of contamination. The previously published method (introduced by (Hill et al., 2011) and refined by (Llobet et al., 2022); see Methods; referred to here as the *Hill-Llobet method*) involves first counting the RP violations then applying a formula that incorporates the total spike count, recording duration, and assumed RP duration to estimate the contamination. The calculation of this estimate relies on the assumption that the base neuron’s spiking rate is uncorrelated with the contamination sources over time (see Discussion for further consideration of this assumption). The estimated contamination is compared to a user-selected threshold corresponding to the maximum level of contamination that is acceptable for the user’s scientific purposes, the *contamination threshold*. Critically, the RP duration is a parameter that must be specified by the user and assumed to reflect the true RP duration of the base neuron accurately.

In the Sliding RP metric, we compute the estimated contamination for each of a range of possible RP duration parameters. A unit with truly low contamination should register low estimated contamination for all values of RP duration up to the base neuron’s true RP duration, then have higher estimated contamination for all larger values, whereas a unit with truly high contamination should register high contamination for every value of RP duration. Accordingly, a unit passes the Sliding RP test when it appears uncontaminated for any tested RP duration. Moreover, rather than computing a point estimate of the contamination, we account for the Poisson spiking statistics to estimate a probability that the unit truly had contamination less than the desired limit. This calculation further improves on prior work by allowing the user to set an interpretable level of confidence that must be obtained in order for a unit to pass the test. The confidence corresponds to the complement probability that the observed violations could have arisen from a truly contaminated unit, i.e. high confidence corresponds to a low probability that the observed violations were generated by an unacceptably contaminated unit. Next, we step through the algorithm in more detail with illustrations of the computation (Fig. 2); see Methods for equations.

The algorithm iterates over a range of possible RP durations, **τ**_r_, that the base neuron might have. In our implementation, **τ**_r_ is tested over the range of 0.5 to 10 ms. For each **τ**_r_, the RP violations are counted (Fig. 2a-c). Rather than using these to compute a point estimate of the contamination level they reflect, the observed count of RP violations (V_o_) is compared to the distribution of counts that would be expected if the unit had the maximum acceptable level of contamination, as specified by the contamination threshold. In this example, the user has specified a contamination threshold of 10%, so the distribution of possible RP violations assuming 10% contamination is derived, given this unit’s total spike count and recording duration (Fig. 2d-f). This distribution is used to determine the probability of having observed V_o_ violations if the unit truly did have 10% contamination. In this example, when **τ**_r_=1 ms, 75 violations were observed, and a unit with 10% contamination is predicted to have 75 or fewer violations 20.2% of the time (Fig. 2d). Thus, for this **τ**_r_, the possibility that the true contamination level was 10% or greater cannot be ruled out with high confidence. We define *confidence* as the complement of the probability that the observed violations arose from a unit with unacceptably high contamination. In this case, then, the confidence is 79.8% at **τ**_r_=1 ms. However, performing the same analysis at **τ**_r_=1.5 ms (Fig. 2b, e) yields a confidence of 91.0%, i.e., there is only a 9.0% chance that this number of violations would have arisen from a unit with the maximum acceptable level of contamination. And, at **τ**_r_=2.2 ms, the confidence rises to 96.9% (Fig. 2c,f). The user sets the desired level of confidence that must be obtained to pass the test, the *confidence threshold*. If the user selected 90% as the confidence threshold, then this unit would pass the test, as one could indeed have >90% confidence that the unit had 10% contamination or less.

For the purposes of illustration, we can compute this same quantity (confidence that contamination is less than the contamination threshold) for a range of possible contamination thresholds that the user might set, depending on their scientific goals. If the user specified a maximum acceptable contamination of 15%, the algorithm would have returned 100% confidence that contamination was <15% across a range of tested **τ**_r_ values, while if the user specified a contamination threshold of 7.5%, the metric would have returned low confidence in contamination being that low across all tested **τ**_r_ values (Fig. 2g). A matrix of confidence values, across tested RP durations **τ**_r_ and across contamination levels, can be computed (Fig. 2h). An iso-confidence line is added at the user-selected 90% confidence threshold (black line), enabling easy visualization of the key results: for maximum acceptable contamination of 10% (red thick horizontal line), the metric does indeed exceed 90% confidence, with a maximum confidence achieved at about 2.5 ms. Moreover, the lowest contamination that can be confirmed with the specified 90% confidence threshold is just under 10%. The algorithm therefore returns both of these values: the highest confidence that can be obtained for contamination being below the user-defined contamination threshold and the lowest contamination that can be confirmed with the user-defined confidence threshold. Either of these are suitable for determining whether a unit passes or fails the test. Thus, the Sliding RP metric is similar in logic to the Hill-Llobet method but tests a range of possible RP durations and considers the probability distribution of violations that may be observed under different hypothetical contamination levels. In this way, the method eliminates the need for the user to select and assume a single RP duration, which may be difficult to do *a priori*. The method meanwhile adds the selection of a desired confidence threshold, a scientifically interpretable parameter reflecting the probability that a unit may have unacceptably high contamination.

### Validation of the metric with simulated data

To validate the metric and compare it quantitatively to previous methods, we simulated spike trains with Poisson-like occurrence and varying values of four parameters: firing rate, RP duration, level of contamination, and recording duration. We applied our metric to these simulated contaminated spike trains, evaluating whether they pass or fail when selecting a maximum acceptable contamination level of 10% and a required confidence level of 90%. For each combination of the simulation parameters, we ran 1000 iterations (Fig. 3, S2).

The metric successfully differentiated between spike trains with high simulated contamination and those without. With sufficiently high recording duration, refractory period, and firing rate – which all contribute to having more spikes and thus greater statistical power – the method sensitively rejected simulated spike trains for which the true contamination was just above the threshold and passed those with contamination just below (Fig. 3a; for the 10 sp/s curve, all simulated spike trains with 8% contamination were correctly accepted, while all simulated spike trains with 12% contamination were correctly rejected). With lower firing rates, the statistical power decreased and the sensitivity (slope) of the rejection curve declined, as expected. At sufficiently low firing rates (0.5 sp/s, for a 2 hr recording duration and 3 ms true RP duration), all simulated spike trains were rejected regardless of contamination (Fig. 3a, dark line along the x-axis). This occurs because even the result of observing zero contaminating spikes will arise with high probability from a spike train with 10% contamination at that firing rate. Thus, observing even zero contaminating spikes does not lend sufficiently high confidence to the conclusion that this spike train had less than the acceptable level of contamination, and all simulated spike trains are accordingly rejected. This is a desirable property of the algorithm, from a conservative perspective, as it ensures that users do not incorrectly take the lack of observed contaminating spikes to support the notion that a unit lacks contamination when this conclusion cannot be made with high confidence (see below, and Discussion, for further consideration of this property).

The Sliding RP metric exhibited two major advantageous features relative to the prior approach (Hill-Llobet method; Fig 3c-d). First, the Hill-Llobet method performed poorly at low firing rates, frequently passing those simulated spike trains with true contamination much greater than the user-defined maximum acceptable limit (Fig 3c, red arrow). This occurs because the Hill-Llobet method incorporates only a point estimate of the contamination rate, rather than accounting for the probabilistic distribution of observed violations that are expected from Poisson processes. In particular, observing zero contaminating spikes will always result in a point estimate of 0% contamination in the Hill-Llobet method, so such spike trains will always pass the test; however, this result becomes increasingly likely to occur even for high contamination levels when firing rates are low. Thus, the Sliding RP method, in accounting for the probability distribution of observed spikes, keeps the false-passing rate approximately fixed to a low level as specified by the user-defined confidence threshold across the entire range of firing rates (but note that the false-passing rate is not perfectly controlled; see Methods section “False passing rate in the Sliding RP metric”, Fig. S3). By contrast, in the Hill-Llobet method the false-passing rate grows arbitrarily large as firing rates decrease. The Sliding RP metric’s correct behavior in limiting false-passing rates according to the user-specified confidence threshold parameter comes at the cost of more conservatively rejecting spike trains that were simulated with true contamination levels below the threshold. This tradeoff is explored further below.

Second, as designed, the Sliding RP metric performed successfully when the true simulated RP durations ranged from 1.5 to 6 ms, while the Hill-Llobet method failed catastrophically when true RP durations were lower than the user-specified threshold (Fig. 3d, red arrow). Spike trains simulated with RP durations of 1.5 or 2 ms, values that were commonly observed in neural data described above, never passed the Hill-Llobet method when used with a RP duration parameter of 3 ms, since ISIs shorter than 3 ms, which occur in these spike trains even when uncontaminated, are considered to reflect contamination. Moreover, since the Sliding RP metric can consider all data up to the true RP duration, the rejection curve becomes more sensitive for spike trains that truly have longer RP durations (Fig. 3b), whereas the Hill-Llobet method ignores any data beyond the selected RP duration.

### Influence of Confidence parameter

The “confidence” parameter of the Sliding RP algorithm allows the user to approximately control the false positive rate of the algorithm, i.e. the proportion of units that pass the test when truly contaminated at or above the acceptable limit. There is no equivalent parameter in the Hill-Llobet approach. When considering simulations with different firing rates, i.e. with different statistical power, the false positive rate scales inversely when using the Hill-Llobet method while remaining approximately fixed, and controlled by the confidence parameter, with the Sliding RP method (Fig. 4a-f). At relatively high firing rates (5 sp/s, Fig. 4a), the performance of both algorithms is overall good, accepting most spike trains simulated with 5% contamination and rejecting most with 15%. But the tradeoff between false passing rate and false rejection rate around the 10% contamination threshold differs. If we consider that spike trains simulated with 12% contamination should all be rejected, so that any passing at that level are defined as false passing, and that spike trains simulated at 8% contamination should all be accepted, so that the percentage passing at that level is the true passing rate, then we can plot a receiver operating characteristic (ROC) curve for the algorithm (Fig. 4d). This visualization illustrates that the confidence parameter adjusts algorithm behavior along the limit of performance (limited by poisson spiking variability), and that the Hill-Llobet approach sits at only one place on this curve, near the Sliding RP method with 80% confidence setting. When the noise is increased, by decreasing the firing rate from 5 sp/s to 2 sp/s, the acceptance curves get less steep (Fig. 4b) and the ROC curve comes closer to the line of unity (Fig. 4e). However, notably, the false positive rates at each confidence setting remain approximately unchanged with the Sliding RP method (ranging from ∼10% to ∼50% as expected), but the Hill-Llobet method has an increased false positive rate, now roughly matching the 70% confidence setting. Finally, with even lower firing rates (1 sp/s; Fig. 4c, f), the Hill-Llobet method’s false positive rate has increased still further and now roughly matches the 50% confidence level. Thus, the confidence parameter allows the user to control the conservativeness of the algorithm in a predictable way.

**Figure 4.**
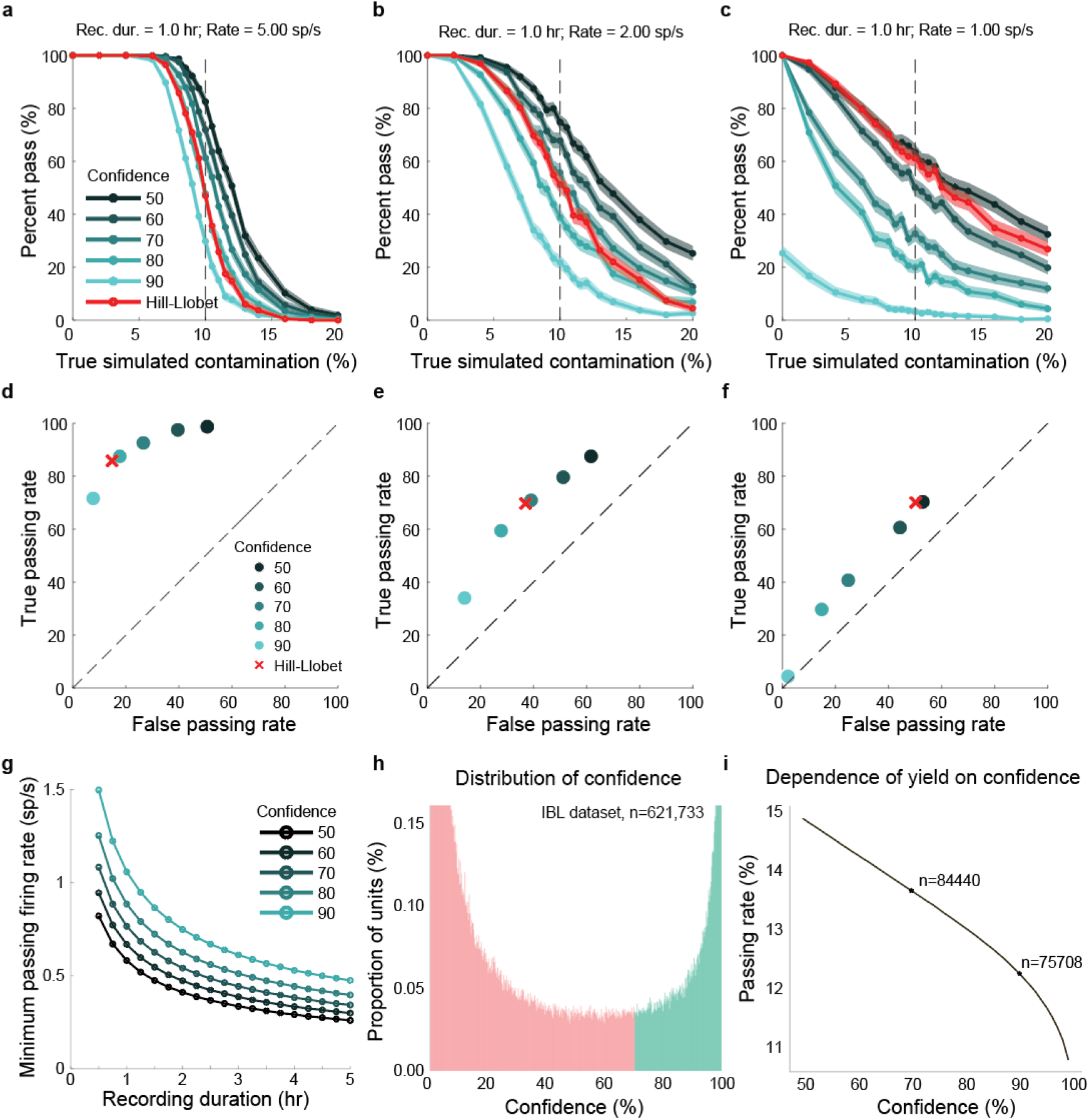
The confidence parameter controls the false passing rate. **a**, Percentage of simulated spike trains passing as a function of true contamination, for varying confidence thresholds (teal shades) and for the Hill-Llobet method (red). Simulated with true RP duration 3 ms; other parameters in plot title. n=1000 simulations per condition; shaded regions reflect 95% confidence intervals. **b**, Same as a, at 2 sp/s. **c**, Same as a, at 1 sp/s. **d**, ROC-style analysis corresponding to a. True passing rate (percentage passing at 8% true contamination) plotted against false passing rate (percentage passing at 12% true contamination) for each confidence threshold. The Hill-Llobet method (red x) occupies a single point on this curve. Dashed line indicates unity. **e**, Same as d, corresponding to b. **f**, Same as d, corresponding to c. **g**, Minimum firing rate at which a simulated spike train with zero contamination can pass the test, as a function of recording duration, for each confidence threshold. Simulated with RP duration 3 ms and contamination threshold 10%. **h**, Distribution of confidence values returned by the Sliding RP metric applied to 621,733 units from the IBL dataset. Units with confidence above 70% (green) pass at that threshold; those below (pink) fail. **i**, Proportion of units passing as a function of the confidence threshold, applied to the same IBL dataset. Total unit counts at selected thresholds are indicated.

One significant consequence of this behavior is that the Sliding RP algorithm will always reject units when their firing rates are too low (for a given recording duration) since accepting a unit with even zero observed RP violations will be inconsistent with the user’s specified confidence (i.e. desired false positive rate). The minimum rate that units need to pass depends strongly on recording duration (Fig. 4g). Note that the exact numbers for minimum firing rate depend on two parameters of the simulation: the true simulated RP duration (here, 3 ms) and the contamination threshold (here, 10%). The relatively high minimum firing rates, particularly for short duration recordings, emphasize two important takeaways from these simulations. First, while it is common for papers to analyze units that have firing rates in the 0.1 - 0.5 sp/s range with recording durations of 1 or 2 hours, it is impossible to be confident that these units are uncontaminated by examining RP violations. Second, the statistical confidence that can be gained in inferences about contamination is significantly improved by recording for longer durations where possible.

Finally, while in general it may seem desirable to use a high confidence threshold (say, 90%) to ensure that few false positives (contaminated units incorrectly accepted by the algorithm) are included for analysis, this parameter does have a small but significant effect on the yield of the algorithm when applied to real data. When applying the algorithm to a dataset of 621,733 units recorded by the International Brain Laboratory (International Brain Lab et al., 2025), changing the confidence threshold from 90% to 70% resulted in 8,732 more units passing (75,708 at 90%; 84,440 at 70%; 11.5% increase; Fig. 4h,i). This difference amounts to about 1.5% of the total number of units. The confidence parameter has the largest effects on spike trains with true contamination levels near to the user-specified threshold, which were few in the IBL dataset. The distribution of these contamination values in other datasets is likely to be dependent on the characteristics of the spike sorting algorithm, and unlikely to be uniform. Hypothetically, if all recorded spike trains had contamination of either 0% or 50%, then the primary effect of reducing the confidence parameter would be to allow more low firing rate units to pass (Fig. 4g).

## Discussion

Here we introduced a spike sorting quality metric, the *Sliding RP metric*, suitable for detecting contaminated units from refractory period violations. Our metric is applicable across brain regions and species, and in individual populations that contain a diversity of neuron types, without parameter tuning. We validated our metric with simulations, demonstrating that it is sensitive, robust, and improves upon performance and interpretability of prior methods. We provide fast, user-friendly code in both MATLAB and Python, including implementation in SpikeInterface (Buccino et al., 2020), to compute the metric from spike trains. This work enables unsupervised and accurate determination of a key aspect of spike sorting quality from electrophysiological datasets.

Our method has several limitations. First, the metric only attempts to quantify the false positive rate of a spike train on the basis of refractory period violations. Since it does not quantify a false negative rate (potential missing spikes), and does not consider other factors that may indicate poor spike sorting, such as inappropriate waveform shapes or amplitudes, it is inadvisable to use it in isolation. Instead, multiple methods may be used in combination to reject units with different types of spike sorting errors. Future work may seek to design methods that detect these other kinds of errors in ways that are also robust to biological variation across brain regions and species.

Second, our method assumes that spike trains are uncorrelated in their rate over time. In most brain regions and in most situations, nearby neurons are weakly but positively correlated on timescales from seconds to minutes (Cohen and Kohn, 2011). In that case, our method is conservative, as these positive correlations may lead to rejecting some units that truly had acceptable contamination levels. To see this, consider that a spike from a contaminating neuron is more likely to land within the RP duration of the base neuron when it is preferentially generated around the time that the base neuron is spiking. This situation has been modeled previously and shown to significantly affect the relationship between observed violations and contamination (Vincent and Economo, 2024). A future extension of our work could attempt to model this correlation and account for it, in order to be less unnecessarily conservative. However, note also that the opposite condition is possible and is more problematic: if the base neuron is negatively correlated with a contaminating neuron, then these two neurons may truly never fire at the same time, and no RP violations will be observed despite potentially significant contamination. This type of situation could occur, for example, with hippocampal place cells, which are known to be minimally active or even silent except in their preferred spatial locations. Thus, units that pass this test must still be interpreted scientifically with caution. Note that this caveat applies equally to the Hill-Llobet method.

Third, our test rejects units with low firing rates even when they have no observed RP violations, in the case that this observation cannot by itself rule out the possibility of unacceptable contamination. This is not a failure or inaccuracy of the method, but rather reveals what was always true from a statistical perspective but has not been widely appreciated or clearly quantified with prior approaches. On the one hand, this is a desirable behavior in that it accurately reflects how an acceptable level of contamination cannot be established for these units on the basis of this method. On the other hand, some users may feel this is unnecessarily harsh: if not a single spike of evidence exists to suggest contamination, perhaps it is fair to assume that the unit is uncontaminated. In that case, users may choose to accept units that have zero RP violations within a chosen RP duration threshold despite failing the test (but, they will have to choose such a threshold arbitrarily or on the basis of prior knowledge, as in earlier approaches). In this case, users should at least be aware that, according to the calculations employed here, they cannot statistically reject the hypothesis that these units are contaminated more than their maximum desired limit, and should interpret their data with appropriate caution. In the future, it may be possible to refine a method such as this by incorporating other indicators of unit quality (such as waveform amplitude) to determine an estimate of the prior probability that a unit is contaminated, which, if available, could be incorporated into the calculation straightforwardly.

Determining the estimated contamination from RP violations has been a bedrock quality control method for extracellular recordings as it can quantitatively, and with rigorous biological grounding, estimate a key source of error that is common in these datasets. Accordingly, essentially every study employing extracellular recordings uses some version of this quality control metric, unless they rely on fully manual - and irreproducible - judgments. The Sliding RP metric introduced here now improves substantially on key aspects of this foundational method. As electrophysiological recordings grow in scale and span an increasing diversity of brain regions and species, the need for quality metrics that are both principled and general will only intensify. We expect that the Sliding RP metric, by removing the need for prior knowledge of refractory period durations and by making explicit the statistical confidence associated with contamination estimates, will be a useful tool for analyzing these datasets.

## Methods

### Macaque dataset for RP duration comparison

All protocols conformed to the guidelines provided by the US National Institutes of Health and the University of Washington Animal Care and Use Committee. Data were collected from three adult rhesus macaques (*Macaca mulatta*). Monkeys were surgically implanted with a titanium headpost and a recording chamber either angled towards the LGN or approximately surface normal to opercular V1.

In the LGN, we recorded extracellular action potentials using Neuropixels NHP 1.0 Long (45 mm; IMEC). The probes were held using a custom adapter that maintained alignment between the probe shank, a guide tube (25 mm-long 20G hypodermic needle), and microdrive assembly (Narishige). Before experiments began, the guide tube was lowered through the dura mater using a thumb screw. The probe was then advanced using a hydraulic microdrive until it penetrated the LGN. Neural activity was verified as originating from the LGN using a hand-held flickering red light.

V1 recordings were conducted similarly using Neuropixels NHP 1.0 Short (10 mm; IMEC). However, a penetrating guide tube was not used to avoid damaging the opercular surface. Instead a flat metal foot was pushed against the dura mater while the probe was advanced through a hole in the foot. The resulting tension alone allowed for the probe to penetrate the dura mater (Namima et al., 2024).

For both regions, the action potential band (sampled at 30 kHz) and local field potential band (sampled at 2.5 kHz) were recorded using OpenEphys (Siegle et al., 2017). For LGN recordings, action potentials were sorted offline using a custom spike sorting pipeline that combined MEDiCINe, a motion correction algorithm (Watters et al., 2025), and Kilosort4 (Pachitariu et al., 2024). For V1 recordings, Kilosort4 was used for both motion correction and spike sorting.

Data were collected while the animals were headfixed and facing a screen. They either fixated or saccaded to targets. Stimuli presented include white noise checkerboard stimuli and drifting gratings. The timing and nature of stimulus presentations was disregarded for the analyses in this work.

### Estimating RP duration for comparison across regions and species (Fig. 1)

To estimate the refractory period of each unit, we fit a sigmoid function to its autocorrelogram (ACG), with steps described in detail below. Note that, while we determined that this procedure was more robust than a threshold crossing or other simple algorithm (due to the complexity, variety, and noisiness of possible ACG shapes), there are nevertheless edge cases that were not captured well by this approach. The inability of any straightforward algorithm to estimate RPs with perfect reliability is what rules out an alternate approach to solving the problem raised in this manuscript (namely, it rules out an approach in which the RP duration is first estimated and then an algorithm to estimate contamination with a fixed RP duration is employed).

First, the ACG was calculated as the count of spikes occurring at each time offset from another spike of the same unit, with bin size of.067 ms (two samples) and window from 0 to 10 ms. We smoothed each ACG using a median filter with a time window of 0.83 milliseconds to reduce noise while preserving the shape of the recovery curve.

Second, to determine the appropriate range for sigmoid fitting, we measured two key points:

Minimum point: We defined the start of the recovery period as the maximum value of the filtered ACG within a narrow time window of 0-0.5 milliseconds. This represents the lowest point of the refractory period. Note that we use the maximum value within this early window (rather than the global minimum) to ensure that subsequent peak detection occurs outside this region.

Maximum point: We detected all peaks in the filtered ACG and applied two criteria to identify valid peaks: (1) the peak value must exceed a threshold defined as 10% of the ACG’s full range (maximum minus minimum), and (2) the peak value must be strictly greater than the minimum point defined above. If no peaks met these criteria, the refractory period estimation was aborted for that unit. Among valid peaks, we selected the one closest to time zero, representing the earliest point of recovery to baseline firing.

Third, we fit a sigmoid function. We truncated the filtered ACG to the interval between the minimum and maximum points identified above, and fit a four-parameter sigmoid function to this segment:

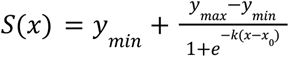

where x is time and there were four fit parameters: y_min_ is the lower asymptote (firing rate during refractory period), y_max_ is the upper asymptote (baseline firing rate), k is the steepness of the curve, and x_0_ is the inflection point (midpoint of recovery). If the fitting procedure failed to converge or produced errors, the refractory period estimation was aborted for that unit.

Finally, we estimated the RP duration from the fit. Once the sigmoid was successfully fitted, we estimated the RP duration as the time point at which the unit has recovered to 10% of the baseline firing:

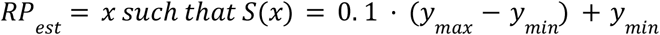

As a final correction, if the estimated refractory period was smaller than the time of the first ACG bin with non-zero firing rate, we set the refractory period to equal the time of that bin instead.

A minority of units had RP_est_ of less than 1 ms, and by visual inspection these were all likely noisy fits and were excluded from quantification for all datasets. For mouse datasets, we selected only units with at least 2 sp/s in total spike rate for inclusion. For macaque datasets, we selected units that had at least 5 times higher firing rate after the estimated RP duration than before.

### The prior state of the art methods for estimating contamination (Hill, Llobet)

The prior state of the art method was introduced in its modern form by Hill et al. (Hill et al., 2011), and based on calculations given by Meunier et al. (Meunier et al., 2003). This calculation estimates the number of RP violations (i.e. instances of an inter-spike interval too small to have arisen from a single neuron) that are expected when exactly two neurons are recorded simultaneously in a single spike train, and each neuron is assumed to be noise-free, i.e. has no RP violations with itself. In this case, rearranging from Hill, the expected number of observed violations is given as:

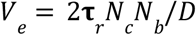

Where V_e_ is the expected number of violations for RP duration **τ**_r_, recording duration D, a count of contaminating spikes N_c_ and a count of spikes in the base neuron N_b_. Since the contamination proportion, C, is given as N_c_/N_t_ where N_t_ is the total number of spikes (i.e. N_t_=N_c_+N_b_), we can re-write this expression as a quadratic in terms of the known variable N_t_:

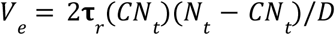

We can then solve for C as a function of the observed number of violations:

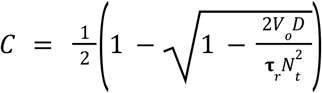

Where V_o_ is the observed number of violations at RP duration **τ**_r_. Either of these equations can be used to determine passing or failure, either by comparing V_e_ at the user-defined threshold contamination level to V_o_, or by comparing the estimated contamination C directly to the threshold.

Though the Hill method has been used extensively, including by us, it arguably does not model the most appropriate situation, in which contaminating spikes arise not from one single other neuron, but instead may arise from multiple other neurons or from a pure noise source, such as from thermal noise fluctuations. In that case, the contaminating spikes can also produce RP violations with each other (i.e. two contaminating spikes may occur within an RP duration from each other), rather than only with spikes of the base unit. The equations describing this situation were derived by Llobet et al. (Llobet et al., 2022) and are given as:

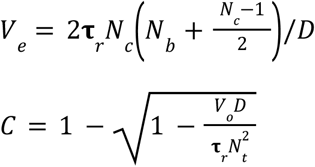

As before, passing and failure can be determined from either expression. We used this calculation in main text figures as the “Hill-Llobet” calculation.

### The Sliding RP metric

The Sliding RP metric estimates the contamination level of a spike train without requiring prior knowledge of the base neuron’s refractory period (RP) duration. The method evaluates confidence that contamination is below a user-specified threshold across a range of possible RP durations, allowing units to pass the test if they appear uncontaminated for at least one tested RP duration (see section below on “False Acceptance Rate of the Sliding RP Metric” for further discussion of this choice of criterion). The *confidence*, as described further below, is defined as the complement probability of having obtained the observed number of RP violations if the true contamination level were at the maximum acceptable level.

#### Input Parameters

The algorithm requires as input:

- S: A vector of spike times in seconds
D: Recording duration in seconds (recommended to specify explicitly, or defaults to maximum spike time if not provided)
- C_thresh_: Maximum acceptable contamination proportion (default: 0.10, i.e., 10%)
- γ_thresh_: Minimum confidence required to accept a unit (default: 90%)
- **τ**_min_: Minimum RP duration to consider in pass/fail decisions (default: 0.0005 s, i.e., 0.5 ms). This value should not be larger than any true RP durations, but selecting a value too small can result in undue contributions of noise. The default value of 0.5 ms should be appropriate for all situations we have considered.

#### Algorithm Description

##### Step 1: Autocorrelogram Computation

The autocorrelogram (ACG) is computed to quantify the number of spike pairs occurring at each inter-spike interval. We compute the ACG by calculating a histogram of differences between all pairs of spike times, with bin size 1/30000 s (the resolution of the recording, for Neuropixels recordings) and range 0 to 10 ms. The resulting ACG, denoted n_ACG_(**τ**), represents the count of spike pairs separated by time lag **τ**.

##### Step 2: Computing Expected Violations

For each combination of tested RP duration (**τ**_r_) and contamination level (C), we compute the expected number of RP violations under the hypothesis that the spike train contains exactly that level of contamination. Following Llobet et al. (2022), we model the situation where contaminating spikes may arise from multiple sources or noise, such that contaminating spikes can produce RP violations with each other in addition to violations with spikes from the base neuron. Given a total spike count N_t_ and contamination proportion C, we define the number of contaminating spikes N_c_ and number of base neuron spikes N_b_ as:

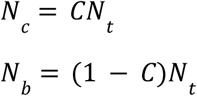

The expected number of RP violations for RP duration **τ**_r_ is:

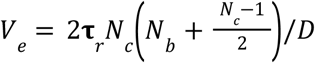

where D is the recording duration. This equation accounts for violations arising from:

1. Contaminating spikes occurring within **τ**_r_ of base neuron spikes: (2**τ**_r_/D) × N_c × N_b
2. Contaminating spikes occurring within **τ**_r_ of each other: (2**τ**_r_/D) × N_c × (N_c - 1)/2

The factor 2**τ**_r_ represents the total window around each spike (**τ**_r_ before and **τ**_r_ after) during which a violation can occur.

##### Step 3: Computing Observed Violations

For each tested RP duration **τ**_r_, we compute the observed number of violations V_o_(**τ**_r_) as the cumulative sum of ACG counts from time 0 up to **τ**_r_:

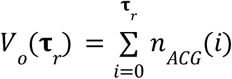

##### Step 4: Computing Confidence Scores

We assume that RP violations follow a Poisson distribution with mean equal to the expected number of violations V_e_. The probability of observing V_o_ or fewer violations, given that the true contamination is C, is given by the Poisson cumulative distribution function:

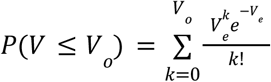

The “confidence score”, γ, that the contamination is actually less than C is defined as the complement of this probability:

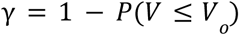

Intuitively, if we observe fewer violations than expected for contamination level C, the confidence that contamination is below C increases. This calculation is performed for all tested contamination levels (by default, 0.5%, 1.0%, 1.5%, …, 35% in steps of 0.5%) and all tested RP durations.

##### Step 5: Pass/Fail Decision

A unit passes the test if there exists any RP duration **τ**_r_ > **τ**_min_ for which the confidence that contamination is below C_thresh_ exceeds γ_thresh_.

The confidence output of the algorithm, γ_max_, is the maximum confidence value across all tested RP durations, at the user-defined maximum acceptable contamination level, C_thresh_.

##### Step 6: Estimating Contamination Level

In addition to the binary pass/fail decision, the algorithm estimates the minimum contamination level C_min_ for which the unit would pass the test at the user-defined confidence level γ_thresh_. If no contamination level meets this criterion, the contamination estimate is set to NaN, indicating high contamination.

Finally, the time of lowest contamination is returned, i.e. the RP duration **τ**_r_ at which the minimum contamination is achieved. This value is one way to estimate the RP duration of the spike train.

##### Outputs

The algorithm returns the following quantities, as defined above:

- P: The pass/fail decision. True when the user-specified maximum contamination (C_thresh_) can be rejected with confidence at least equal to the user-specified minimum (γ_thresh_), and false otherwise.
- γ_max_: The maximum confidence level obtained for the user-specified maximum contamination (C_thresh_)
C_min_: The minimum contamination level that can be confirmed at the user-specified minimum confidence level (γ_thresh_). When none of the tested contamination levels pass at confidence level γ_thresh_ (for the default testing range of up to 35% contamination tested, this would be when the estimated contamination is > 35%), then this return argument will be NaN.
- C_all_: The full confidence matrix of size [n_C_ × n_**τ**_], where n_C_ is the number of contamination levels tested and n_**τ**_ is the number of RP duration bins, for detailed inspection
- C_tested_: Vector of tested contamination levels (as percentages)
- **τ**_tested_: Vector of tested RP durations (in seconds)
- n_ACG_: The autocorrelogram values at each RP duration

##### Computational Considerations

The confidence matrix computation scales as O(n_C_ × n_**τ**_), where n_C_ is the number of contamination levels tested and n_**τ**_ is the number of RP duration bins. For rapid computation when only the pass/fail decision is needed, users may specify the set of contamination levels to test as a single value (equal to C_thresh_), though this precludes accurate estimation of the contamination level.

### False acceptance rate in the Sliding RP metric

The Sliding RP calculation is designed to reject units at the threshold contamination (C_thresh_) with a probability given by the user-defined confidence threshold (γ_thresh_). However, it can be observed that the metric imperfectly controls this probability: in simulated spike trains with γ_thresh_=90%, the percentage of units simulated with the threshold contamination level that pass is around 30% for most parameters (Fig 3a-b) rather than the desired 10%. This arises because the method evaluates the passing criterion across a range of refractory period durations (**τ**_r_), so it fundamentally involves multiple statistical comparisons. Since these comparisons evaluate monotonically expanding windows of the same autocorrelogram, the tests are highly correlated. Standard multiple-comparison corrections (e.g., Bonferroni) assume independent tests and would severely over-penalize the metric, reducing statistical power. To correctly control the family-wise error rate without sacrificing power, we modeled the accumulation of expected violations as a single Markov chain and computed the exact first-passage probability of a Poisson process.

Under the null hypothesis that a unit’s true contamination exactly equals the user-defined threshold C_thresh_, the expected number of violations at time bin k is *V*_*e*_ (**τ**_*k*_). For each time bin, we define a critical boundary *c*_*k*_, representing the maximum number of observed violations that would result in a nominal pointwise confidence score greater than or equal to the target γ_thresh_:

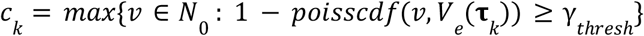

Using a 1D dynamic programming approach, we calculate the probability P_false_ that a randomly generated discrete random walk with independent Poisson increments *λ*_*k*_ = *V*_*e*_ (**τ**_*k*_) − *V*_*e*_ (**τ**_*k*−1_) will dip below the boundary *c*_*k*_ at any point across the entire sliding window. The exact, family-wise corrected confidence score is then given by *γ*_*corrected*_ = 1 − *P*_*false*_. This exact correction successfully locks the false-passing rate to the desired α (where α=1−γ_thresh_) when the full tested time window is subject to the null hypothesis (e.g., for simulated units with long true RP durations; Fig. S3a).

We note that the test becomes strictly conservative for spike trains for which the true RP duration is much shorter than the maximum tested window **τ**_max_. For a unit with short RP duration, actual spikes from the base neuron rapidly accumulate in the autocorrelogram at **τ**>**τ**_true_, making it effectively impossible to randomly cross the boundary late in the analysis window. Because the algorithm corrects for the statistical “chances” to pass across the entire window up to **τ**_max_, this correction represents an over-penalty for these units. Consequently, the empirical false passing rate for short-RP neurons falls well below that expected from the user-defined confidence threshold (Fig. S3b).

Since this correction cannot be exact without knowledge of the true RP duration, and since a method that does not depend on knowledge of the true RP duration was desired, we have omitted it from the standard implementation of the algorithm (and from all plots except Fig. S3). Moreover, this method incurs a significant computational cost. Despite the omission of this approach for multiple-comparisons correction, the false passing rate is nevertheless approximately controllable in a predictable manner (Fig. 4).

### Spike train simulations

Simulations were performed by generating synthetic spike trains as Poisson processes with the given total spike rate, RP duration for the base neuron, recording duration, and contamination proportion. First we calculated the base rate (rate in spikes/s of the base neuron) and contamination rate as:

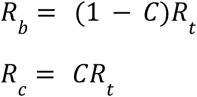

Where R_t_ is the total rate of the spike train, R_b_ is the base rate, R_c_ is the contamination rate, and C is the proportion of contamination. We then generated a vector of randomly-generated interspike intervals, *I*, which are given as:

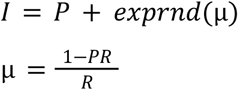

Where P is the RP duration and µ is a corrected rate, accounting for the existence of the RP duration, such that the total desired rate is correct. The function *exprnd* returns an exponentially distributed random number with mean parameter µ, and the expression for *I* is evaluated a sufficient number of times such that the cumulative sum of *I* is greater than the recording duration. The spike train (i.e. vector of spike times in seconds) is obtained as the cumulative sum of *I*. Spike times beyond the recording duration are dropped.

This procedure for generating spike trains is applied separately to generate a spike train for the base neuron using rate R_b_ and RP duration P, and a spike train for the contamination with rate R_c_ and no RP (i.e. P=0). These spike trains are appended and sorted to obtain the final simulated spike train.

A total of 1000 simulations were run with each combination of parameters (firing rate, recording duration, RP duration, contamination) and the proportion of the 1000 that passed was computed. 95% confidence intervals on this proportion were calculated with the Clopper-Pearson method (function ‘binofit’ in Matlab).

## Acknowledgements

We thank A. Ladd and J. Siegle for helpful feedback on this manuscript.

This work was supported by the Simons Foundation Collaboration for the Global Brain (to the IBL), the Wellcome Trust (216324/Z/19/Z to the IBL), National Institutes of Health (U19 NS123716 to NAS and the IBL; R01 EY018849 to GDH; T32 EY007031 and F31EY037124 to LMB; P51 OD010425 to the WaNPRC), the National Science Foundation (CAREER 2142911 to NAS), the Pew Biomedical Scholars Program (to NAS), and the Klingenstein-Simons Fellowship in Neuroscience (to NAS).

## Author Contributions

NAS conceived the method, wrote the MATLAB implementation and simulations, performed data analysis, and wrote the manuscript. NR wrote the initial Python implementation, performed data analysis and simulations. GC and OW reviewed and revised the Python implementation and developed unit testing. OW performed analyses on IBL data (Fig. 4h,i). OW packaged the Python code and developed continuous integration. RAR, LMB, RAC, and GDH performed macaque data collection and preprocessing. All authors reviewed and edited the manuscript.

## Code and Data Availability

Code for the metric and for the simulations presented is available at https://github.com/SteinmetzLab/slidingRefractory/.

## Supplemental Figures

**Supplemental Figure S1.**
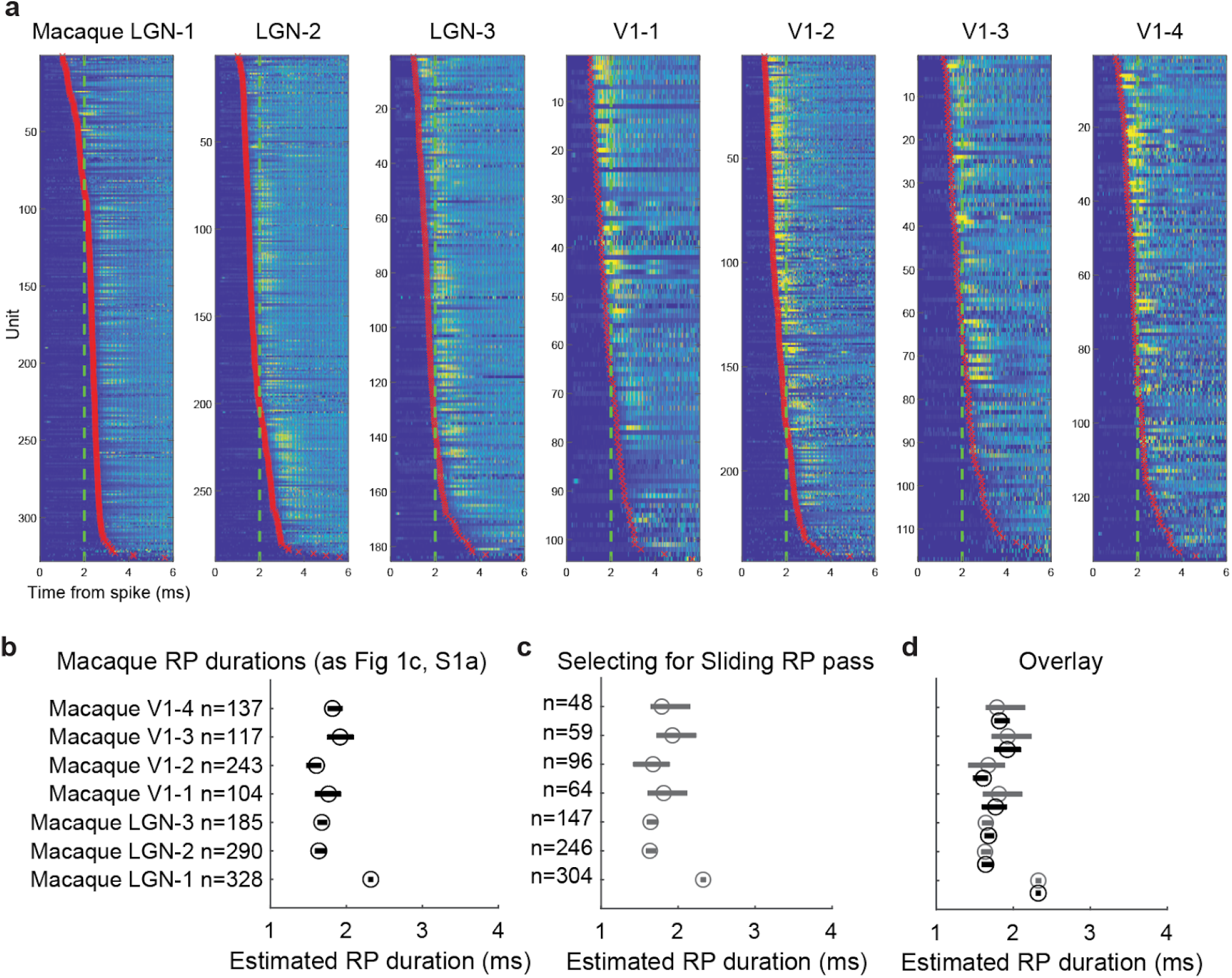
Detail of estimated RP durations for macaque data. **a**, ACGs for all included macaque neurons. Each subplot represents one of the 7 recordings in Figure 1. Each row represents the ACG of one neuron with color value indicating normalized amplitude. Red markers indicate the estimated RP duration from the sigmoid fitting algorithm. Green dashed line corresponds to 2 ms. The first recording (LGN-1) is anomalous in having a median RP duration of > 2 ms and only a few neurons with RP duration < 2 ms. All six of the others have median < 2 ms and many clear examples of RP duration < 2 ms. **b**, Reproduction of median and bootstrapped CI’s of RP durations for each macaque dataset (Fig. 1c). **c**, Same as b, but further selecting only those units that also pass the Sliding RP test. **d**, Overlay of b and c, showing that contaminated units do not notably bias these RP duration estimates.

**Supplemental Figure S2.**
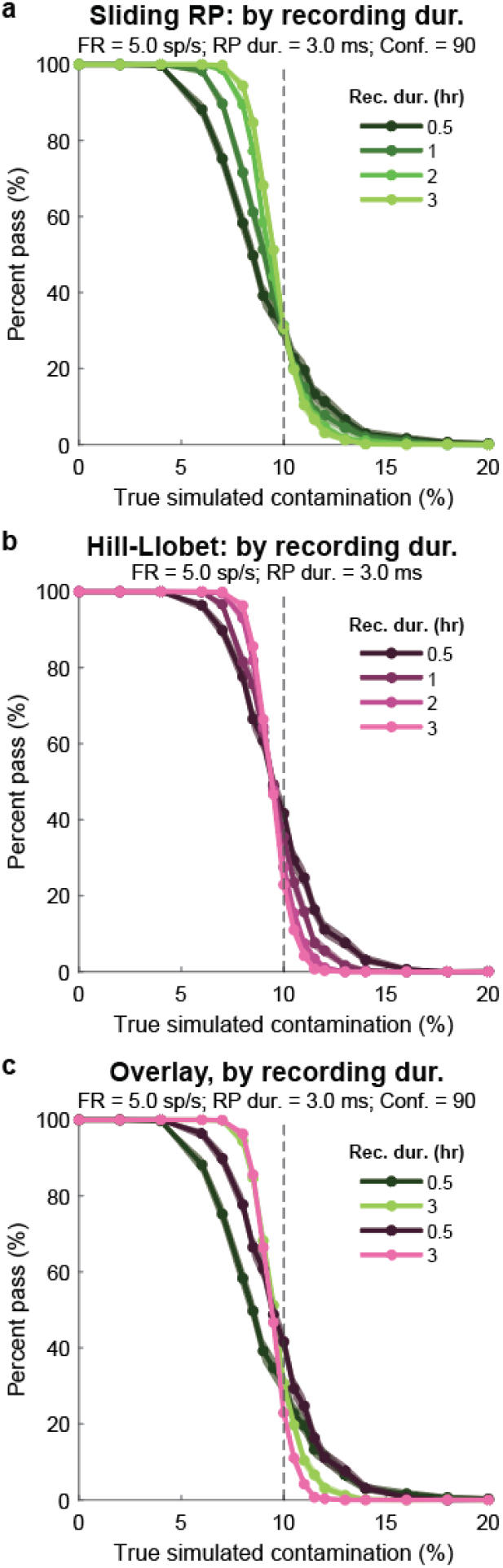
Performance of the metric for different recording durations. **a, b, c**, As in Fig 3 but for different simulated values of recording duration.

**Supplemental Figure S3.**
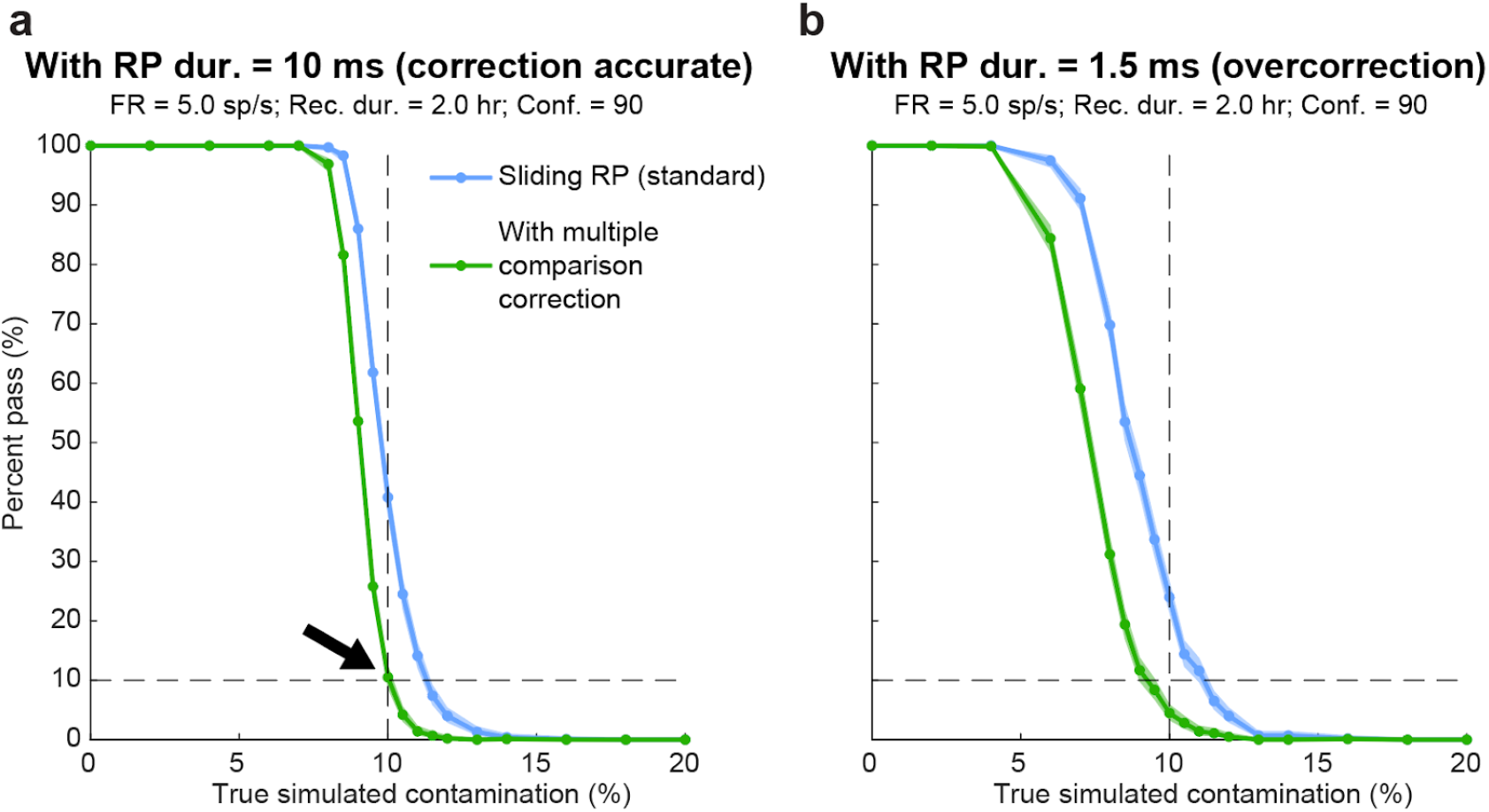
A correction for multiple comparisons accurately controls the false acceptance rate, but only when the RP duration is known. **a,** Percentage of simulated spike trains passing the test as a function of simulated contamination, for the standard Sliding RP test and the modified version with a multiple comparisons correction. In this simulation the RP duration is 10 ms, i.e. matching the window over which the Sliding RP algorithm tests, and matching the assumptions of the multiple comparison correction. Note that the correction works perfectly: the green trace returns exactly 10% passing at 10% contamination (black arrow; for confidence threshold = 90 and contamination threshold = 10%). **b**, As in a but for a simulated RP duration of 1.5 ms. When the true RP duration is shorter than the tested window, the multiple comparisons test now results in an overcorrection (only ∼5% pass at 10% simulated contamination). The requirement of this correction to know the RP duration in order to perform appropriately is precisely what was to be avoided in the design of the Sliding RP method, so this approach is not preferred.

